# AetherXeno: an allele-resolved receptor-occupancy atlas for regulatory pharmacogenomics

**DOI:** 10.64898/2026.09.20.752952

**Authors:** Yen-Hung Chen, Teng-Che Chuang, Hsiang-Wen Lin, Yi-Chuan Li

## Abstract

**Motivation:** Interpreting noncoding pharmacogenomic variants requires identifying the regulatory program that may be perturbed and the direction of the effect, information not supplied by population frequency or general regulatory annotations.

**Results:** We developed AetherXeno, an allele-resolved atlas that scores all substitutions across 8,691 experimentally supported elements for PXR, FXR and AhR and connects 17.27 million variants to evidence matched by exact allele. Across the six paired comparisons, the gain in Spearman correlation ranged from 0.039 to 0.334. Every regional bootstrap interval excluded zero. Homology exclusion preserved these findings, while motif core and liver expression enrichment supported biological organization of the score landscape. The browser, programmatic interface and bulk files expose the same versioned records from individual variants to cohort analyses.

**Availability and implementation:** The web resource is available at https://aetherxeno.org and the versioned data at doi:10.5281/zenodo.20753827.

## 1 Introduction

Noncoding variation can modify transcription factor occupancy without altering a protein-coding sequence. This presents a specific problem in pharmacogenomics because drug disposition and response depend on regulatory programs controlling enzymes and transporters. Receptors activated by ligands integrate xenobiotic, metabolic and environmental signals. A variant within an element bound by a receptor may therefore increase one regulatory program, decrease another or have little predicted effect. Population frequency, conservation and broad functional scores remain useful, but none alone identifies the affected receptor or predicted direction of occupancy change.

Pregnane X receptor (PXR), farnesoid X receptor (FXR) and aryl hydrocarbon receptor (AhR) provide experimentally tractable examples of this problem. ChIP-seq studies in primary human hepatocytes have mapped receptor binding after exposure to rifampicin, GW4064 and TCDD, respectively (Filipovic et al. 2023; Smith et al. 2014; Zhan et al. 2014). These experiments define regulatory elements relevant to xenobiotic sensing, bile acid signaling and environmental response. Yet their direct reuse for variant interpretation is difficult: the measured signal is stored as genomic coverage or peak intervals, whereas a practical pharmacogenomic query begins with a variant specified by chromosome, position, reference allele and alternative allele and asks how that substitution may alter receptor occupancy.

Sequence models such as Enformer and AlphaGenome predict broad molecular signals (Avsec et al. 2021, 2026). RegulomeDB catalogues regulatory annotations, and MPRAVarDB catalogues experimentally assayed variants (Boyle et al. 2012; Jin et al. 2024). None provides a precomputed map of alternative and reference effects for each receptor in the hepatocyte experiments analysed here. AetherXeno fills this gap by scoring all three substitutions at each supported base and linking observed alleles to external evidence.

Because regulatory evidence is allele specific, each release preserves GRCh38 chromosome, position, reference allele and alternative allele. Predicted effects, external evidence and candidate pharmacogene links remain separate. A reader can therefore trace each hypothesis without treating proximity or a model score as clinical evidence.

AetherXeno adapts a frozen AlphaGenome sequence trunk to three receptor occupancy datasets from primary human hepatocytes. It distributes directional effects through a browser, programmatic access and versioned bulk files. Each allele is linked to population frequency, liver expression, clinical interpretation, trait association, motif and conservation resources. Methods describes the source databases, versions and matching procedures. We tested whether a separate head for each receptor recovers measured

ChIP-seq signal better than native AlphaGenome and Enformer outputs on chromosomes reserved for testing. We repeated the benchmark after homology exclusion and evaluated motif and liver expression enrichment. Published reporter alleles were treated as recovery of variants used during source selection. These analyses assess prioritization of receptor signal. Clinical response prediction was outside the study scope.

## 2 Methods

### 2.1 Study design and source receptor occupancy experiments

The study comprised construction of receptor occupancy targets, adaptation of a prediction head for each receptor, exhaustive in silico saturation mutagenesis and integration of evidence matched by allele. All coordinates were represented on GRCh38. Inputs were public ChIP-seq experiments in primary human hepatocytes, retaining the exposure context of each study.

For PXR, we pooled libraries from dimethyl sulfoxide and rifampicin treatments and used input DNA as control. The target therefore represents constitutive occupancy together with response to ligand rather than an isolated rifampicin response. The FXR experiment followed GW4064 exposure and used matched rabbit immunoglobulin G as background. The AhR experiment followed 1 nM TCDD exposure for 24 h and used input DNA as control (Filipovic et al. 2023; Smith et al. 2014; Zhan et al. 2014). The AhR experiment had one biological replicate. Supplementary Table S1 gives accessions, read layouts and read lengths.

### 2.2 Read processing and definition of receptor elements

Reads were aligned to GRCh38.p13 with BWA-MEM 0.7.19, sorted by coordinate, marked for duplicates and filtered at mapping quality 20 (Li 2013). Peaks were called with MACS2 2.2.9.1 using the control and threshold chosen for each receptor (Zhang *et al*., 2008). Supplementary Table S2 gives the complete parameters.

Because AhR had one replicate, its calls were intersected with ENCODE HepG2 ATAC-seq peaks. Two prespecified canonical target peaks were restored after filtering (ENCODE Project Consortium 2020). Peak summits were ranked, and overlapping or invalid windows were consolidated. Supplementary Methods S2 and Tables S2 and S7 report aggregate counts, counts for each receptor, the genomic union and ENCODE file identifiers.

### 2.3 Adaptation of prediction heads for each receptor

We used AlphaGenome research implementation v0.2.0 and its fold 0 research checkpoint (Avsec et al. 2026). The sequence trunk was frozen, and a separate ChIP-seq prediction head was fitted for each receptor. Inputs were 131,072 bp and outputs had 128 bp resolution. Adam optimization followed Kingma and Ba (2015). Supplementary Table S2 reports learning rates, step counts, gradient clipping and the exact checkpoint identifier.

Five chromosomes were excluded during fitting of receptor heads. Published PXR variants on chromosomes 2 and 20 later informed the choice between the pooled PXR target and a target based only on rifampicin treatment. Although not training labels, they could influence that input choice. The primary analysis therefore used the three unaffected holdouts, chromosomes 8, 11 and 15, without refitting or retuning.

### 2.4 Exhaustive saturation scoring and normalization

We generated and evaluated each possible nonreference allele at every base of the receptor element union. Let *f*_*r*_(*x*)_*j*_ denote the prediction from receptor head *r* in output bin *j*, and let *c* denote the bin containing the queried position. The raw difference between alternative and reference predictions for variant *v* was

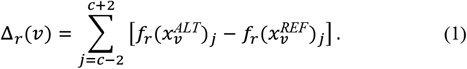

The sum across five bins covers 640 bp centred on the variant. To retain direction while supporting comparison across receptor heads, one denominator was defined separately for each receptor on the complete saturation population *V*:

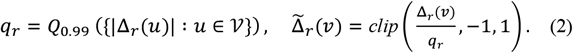

Supplementary Table S7 reports the p99 denominator for each receptor and the neutral display interval. Positive and negative normalized values indicate predicted increases and decreases in occupancy, respectively. The normalized value is not a probability, percentile or pathogenicity score. Variants were ranked across receptors by the unclipped statistic

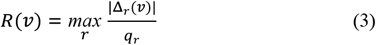

Ties were resolved by stable variant identifier. The exact top 1% was defined after this ordering among substitutions within supported receptor elements. Raw effects, normalized effects and the ranking statistic are retained in the bulk release.

### 2.5 Evidence integration, prioritization strata and pharmacogenomic context

All external evidence was joined by GRCh38 chromosome, position, reference allele and alternative allele. Population observations came from gnomAD v4.1, expression associations from GTEx V10 liver, clinical assertions from ClinVar and trait associations from the NHGRI–EBI GWAS Catalog (Cerezo et al. 2025; Chao et al. 2024; GTEx Consortium 2020; Landrum et al. 2025). Evidence attached to one allele was not propagated to other substitutions at the same position.

JASPAR 2024.1 CORE vertebrate matrices were scanned with FIMO at a nominal threshold of *P* < 1 × 10^−5^ (Grant et al. 2011; Rauluseviciute et al. 2024). The hg38 phyloP track supplied conservation scores across 100 species (Pollard et al. 2010). Supplementary Methods S4 reports the retained matrices, background model and annotation procedure.

Two transparent review strata were defined. Tier A contains GTEx liver expression records matched by exact allele. Tier B contains variants in the top 1% by score with gnomAD allele frequency below 0.01. Both strata are search aids, not pathogenicity or treatment classes.

The curated pharmacogenomic layer connects an element bound by a receptor to a candidate drug disposition gene when a production peak lies within 25 kb of the transcription start site. Each chain retains the receptor, exposure, regulatory locus, predicted consequence, drug context and literature support as separate fields. This structure follows an evidence-based pharmacogenomics framework (Whirl-Carrillo et al. 2021). Before a chain can enter a versioned public release, a pharmacist reviewer assesses the proposed receptor–gene mechanism against the cited evidence. The decision is recorded as accepted or requiring revision; corrected mechanism wording and reviewer notes are retained when revision is required. This review addresses mechanistic pharmacology, not inherited PharmGKB drug lists, clinical actionability, dosing or prescribing advice. Proximity does not establish the target gene. Supplementary Methods S4 and Table S7 report chain counts and construction details.

### 2.6 Web application and versioned release

The web application was built with Next.js and React. It queries a read-only PostgreSQL database hosted on Supabase. Users can search by coordinate, rsID, interval, gene or receptor element and compare receptors. Programmatic endpoints and Parquet files support larger analyses. Pagination and interval limits prevent unbounded responses. The noncommercial service requires no registration. Zenodo record 20753827 archives versioned tables, dictionaries, provenance and checksums (Chen et al. 2026). Each artifact states its source and data terms.

### 2.7 Primary test set and comparative evaluation

Balanced receptor peak and accessible background windows from chromosomes 2, 8, 11, 15 and 20 formed the test set. Published PXR variants on chromosomes 2 and 20 had influenced source selection. The primary test therefore used chromosomes 8, 11 and 15, while the complete set was retained for sensitivity analysis. Supplementary Tables S3 and S4 report exact window and bin counts. No model was refitted.

The same bins were scored by AetherXeno and the two comparators. For AlphaGenome, HepG2 RXRA and AHR outputs served as the available proxies for PXR and FXR and for AhR, respectively. For Enformer, we averaged one liver CAGE output and three liver H3K27ac outputs (Avsec et al. 2021). Measured occupancy came from the processed ChIP-seq signal. We excluded bins overlapping an ENCODE HepG2 H3K27ac track to define the stratum with low baseline activity. Supplementary Table S7 provides accession details.

The primary endpoint was Spearman correlation between predicted and measured signal. Comparative effects were paired differences between AetherXeno and each comparator in identical bins. Regional block bootstrap confidence intervals preserved dependence within each window. Supplementary Table S4 reports exact resample and postfilter denominators. The area under the receiver operating characteristic curve for distinguishing peaks from background was secondary because regional activity confounded this contrast. Correlation measures the ranking of receptor occupancy. It does not measure accuracy for individual variants or clinical outcomes.

### 2.8 Control for sequence homology across chromosomes

Chromosome holdout prevents coordinate overlap but does not exclude duplicated sequence on other chromosomes. We therefore aligned every unique test window, including the primary subset, with minimap2 against a conservative proxy for training data from the remaining chromosomes (Li 2018). Supplementary Methods S7 reports exact window counts and defines the proxy. Because the proxy includes sequence beyond the intervals used to fit receptor heads, it can overestimate homology between training and test sequences.

We evaluated sequences before and after soft masking with Repeat-Masker at four prespecified combinations of alignment length and identity (Supplementary Table S5; Smit et al. 2015). The sensitivity analysis excluded matches that remained after masking and spanned at least 1 kb with 99% identity. Frozen predictions were then reevaluated. This analysis addresses separation between sequences used for fitting and testing, not AlphaGenome pretraining.

### 2.9 Orthogonal biological analyses

For liver expression quantitative trait locus enrichment, variants observed in gnomAD within supported receptor elements were collapsed by genomic position using the maximum absolute score for each receptor. We refer to these expression associations as eQTLs below. Significant GTEx V10 liver eQTL pairs were reduced to the variant with the lowest nominal *P* value for each gene and intersected by position. The top and bottom score quintiles were compared for each receptor and for the maximum across receptors by

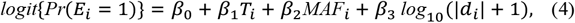

where *E*_*i*_ denotes a lead eQTL position, *T*_*i*_ indicates the top quintile, *MAF*_*i*_ is minor allele frequency and *d*_*i*_ is distance to the nearest transcription start site. The adjusted odds ratio was *exp*(*β*_1_). An ordinal model tested the trend across quintiles. Chromosome was added in sensitivity models when the sparse event design remained estimable. This analysis by genomic position tests enrichment, not the signed direction or causality of an individual eQTL.

Motif enrichment compared substitutions in the top 1% by score with all remaining substitutions in supported receptor elements using a two-sided Fisher’s exact test. Local motif grammar was summarized as the ratio between mean maximum absolute raw effect at each position inside a retained motif core and the corresponding mean in flanks extending ±50 bp. Core and flank distributions were compared using a one-sided Mann– Whitney U test. Motif occurrence and eQTL status were not targets used to train receptor heads.

Published reporter alleles were assessed only as recovery of variants used during source selection. Exact contrasts with experimentally evaluable directions and indeterminate cases were prespecified from the evidence record (Gotoh-Saito et al. 2025; Sugatani et al. 2008) and are listed in Supplementary Table S8B. Because every candidate reporter had informed the choice of source input, the independent accuracy denominator is zero.

### 2.10 Representative use case with exact allele matching

The locus example was selected by an explicit completeness rule. The rule required a released record within the display bounds, with population frequency, a liver expression association, a trait association and elements for more than one receptor. The first eligible record was rs2054576 A>G at a locus linked to *ABCG2*. This record illustrates atlas navigation. It is not an independent evaluation endpoint.

## 3 Results and discussion

### 3.1 A saturation atlas with separate scores for each receptor

Processing of the occupancy datasets from primary human hepatocytes produced 8,691 receptor element records. Exhaustive saturation and integration with population data yielded 17,267,831 unique variant records with separate PXR, FXR and AhR effect fields. Figure 1 and Supplementary Table S7 report the exact saturation, overlap and genomic coverage counts.

**Figure 1.**
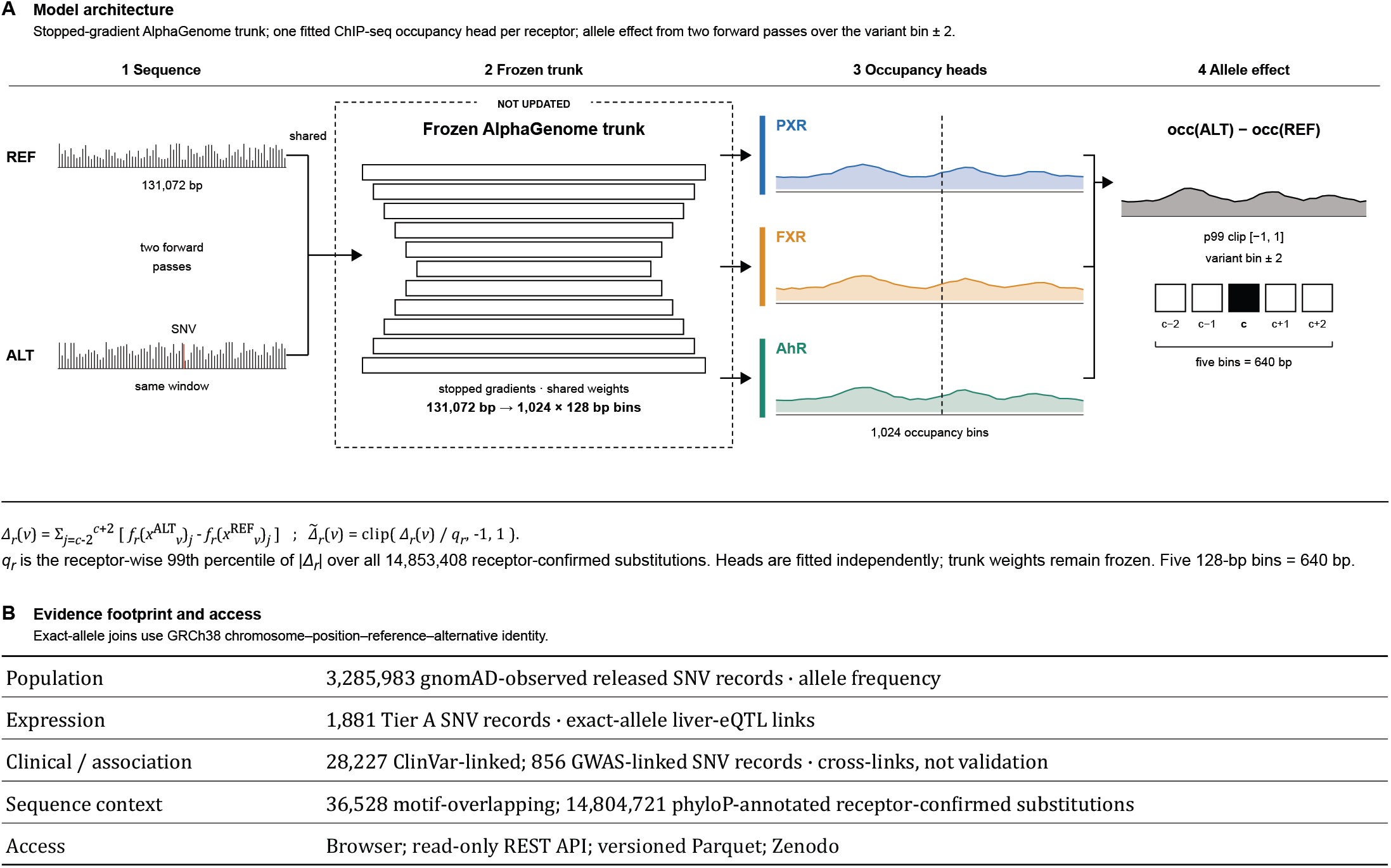
Model architecture and evidence footprint of AetherXeno. (A) Receptor-specific ChIP-seq occupancy heads operate above a frozen AlphaGenome trunk, with signed allele effects derived from matched reference and alternative forward passes. (B) Evidence layers matched by exact allele and access routes.

The release retained raw allele effects, signed values normalized for each receptor and the unclipped ranking statistic. Users can inspect direction and magnitude for one receptor while retaining a deterministic ranking across receptors. Two transparent review strata reduce the search space while leaving the continuous scores available. Supplementary Table S7 reports their component counts.

The atlas also links each released allele to clinical records, trait associations, retained motifs and phyloP conservation. Annotation coverage is summarized in Figure 1 and Supplementary Table S7. Links between receptors and candidate genes are treated as hypotheses. They require pharmacist review before release and are not clinical assertions. The browser, programmatic interface and bulk files use the same identifiers and fields.

### 3.2 Receptor adaptation improves signal ranking in test data

Because PXR variants on chromosomes 2 and 20 informed source selection, the primary benchmark used chromosomes 8, 11 and 15. Supplementary Table S3 reports the sensitivity analysis across all five chromosomes.

Figure 2A shows Spearman correlations of 0.211, 0.432 and 0.638 for PXR, FXR and AhR, respectively. Each receptor exceeded both comparators. Across the six paired comparisons, gains ranged from 0.039 to 0.334 and every regional bootstrap 95% confidence interval excluded zero. Figure 2B and Supplementary Table S4 report the comparisons.

**Figure 2.**
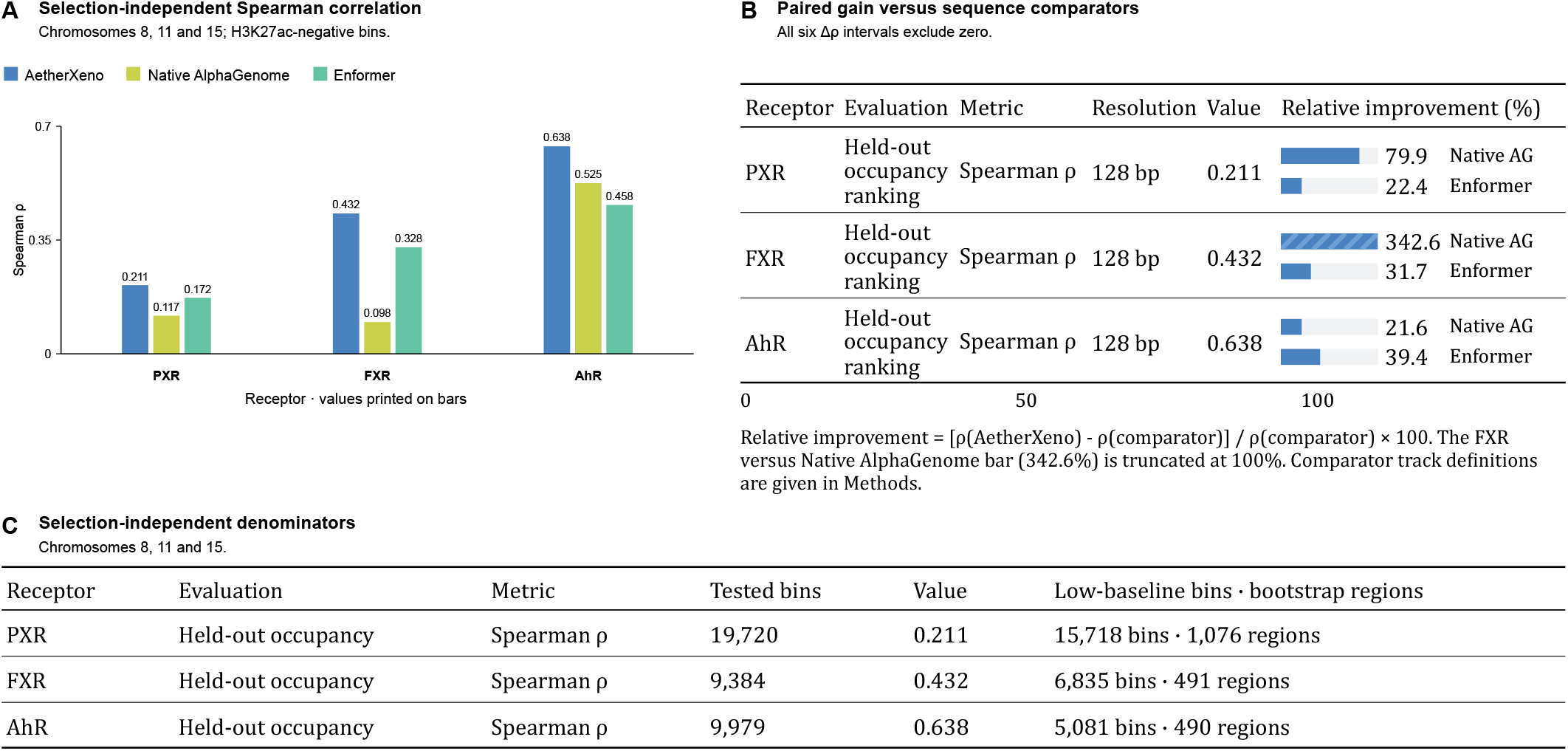
Primary benchmark of receptor signal. (A) Spearman correlations on chromosomes 8, 11 and 15. (B) Relative improvements over Native AlphaGenome and Enformer; the FXR comparison with Native AlphaGenome is truncated at 100%. (C) Denominators for tested bins, bins with low baseline activity and bootstrap regions.

### 3.3 Comparator conclusions persist after homology exclusion

Repeat masking left 18 matches of nearly identical sequence across chromosomes, including four in the primary subset. Supplementary Figure S2 and Table S5 report the screen. Excluding those four regions changed no rounded correlation and left all six comparator intervals above zero. Detectable homology between chromosomes therefore did not explain the primary benchmark.

### 3.4 Interpretation based on exact allele matching at an *ABCG2* locus

Figure 3A and B show the observed rs2054576 A>G allele in overlapping PXR and FXR elements linked to *ABCG2*. Its population frequency is 8.1%. The allele is associated with *ABCG2* expression in liver and with hyperuricemia, as shown in Figure 3C.

**Figure 3.**
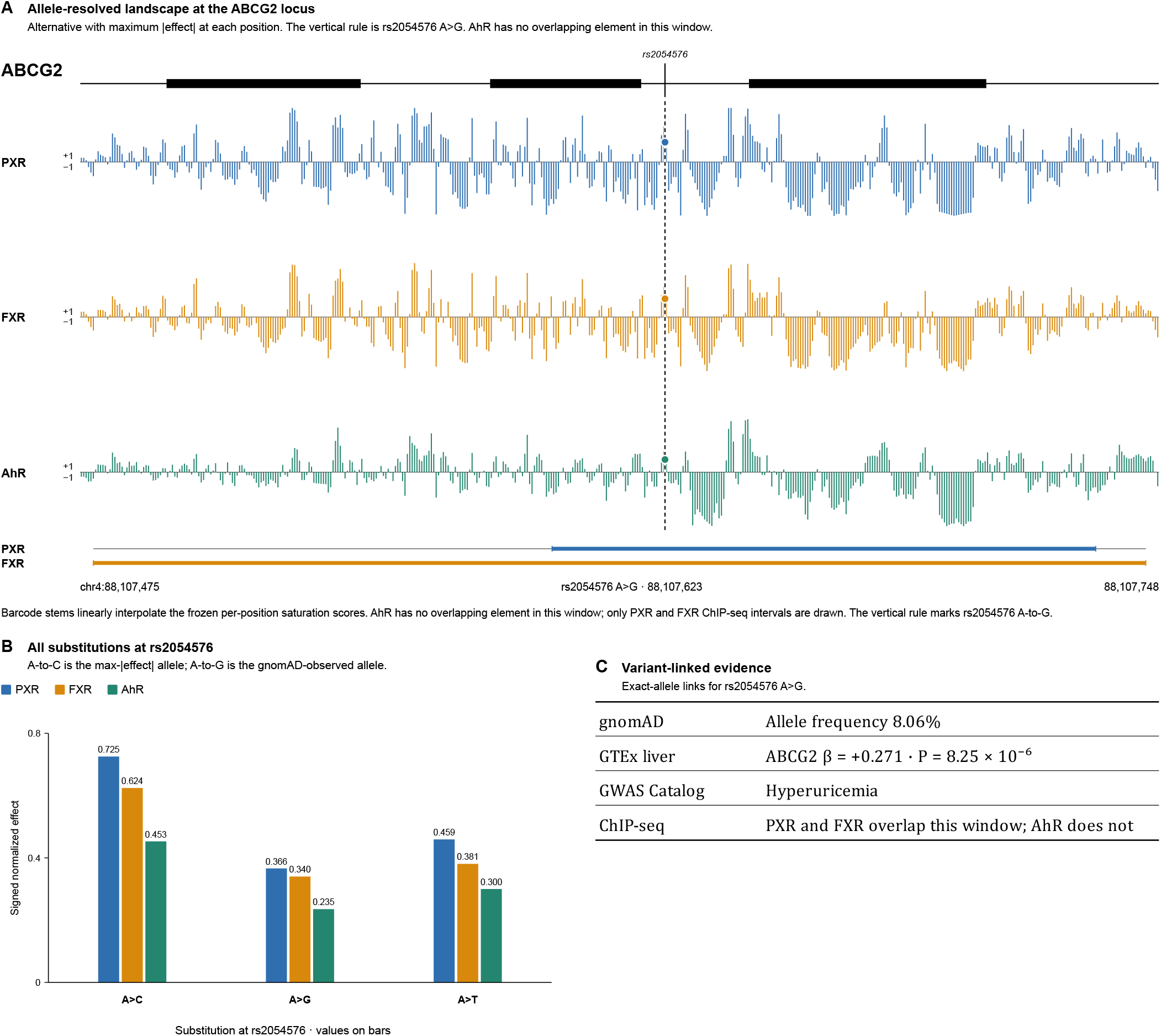
Interpretation based on exact allele matching at the *ABCG2* locus. (A) Receptor effect tracks on shared genomic coordinates and observed A>G scores with supported PXR and FXR intervals. (B) All substitutions at the position with reference allele A. (C) Population, liver eQTL, GWAS and ChIP-seq evidence matched to A>G.

The same record can be found by coordinate, rsID, interval or gene in the browser and retrieved programmatically or from the bulk release. This example shows how AetherXeno moves from prioritization across the genome to evidence review for a specific allele. It nominates a testable hypothesis linking a receptor to a gene.

### 3.5 Orthogonal motif and expression analyses

The motif enrichment odds ratio was 19.62. Mean sensitivity was 1.67 to 2.04 times higher in motif cores than in adjacent flanks across the three receptors. Figure 4A and Supplementary Table S8A report these analyses. Liver eQTLs were also enriched in the highest score quintile after adjustment for allele frequency and distance to the nearest transcription start site. Adjusted odds ratios ranged from 2.63 to 5.10. Figure 4B and Supplementary Table S6 report the expression analysis. Together, these results show biological organization of the score landscape.

**Figure 4.**
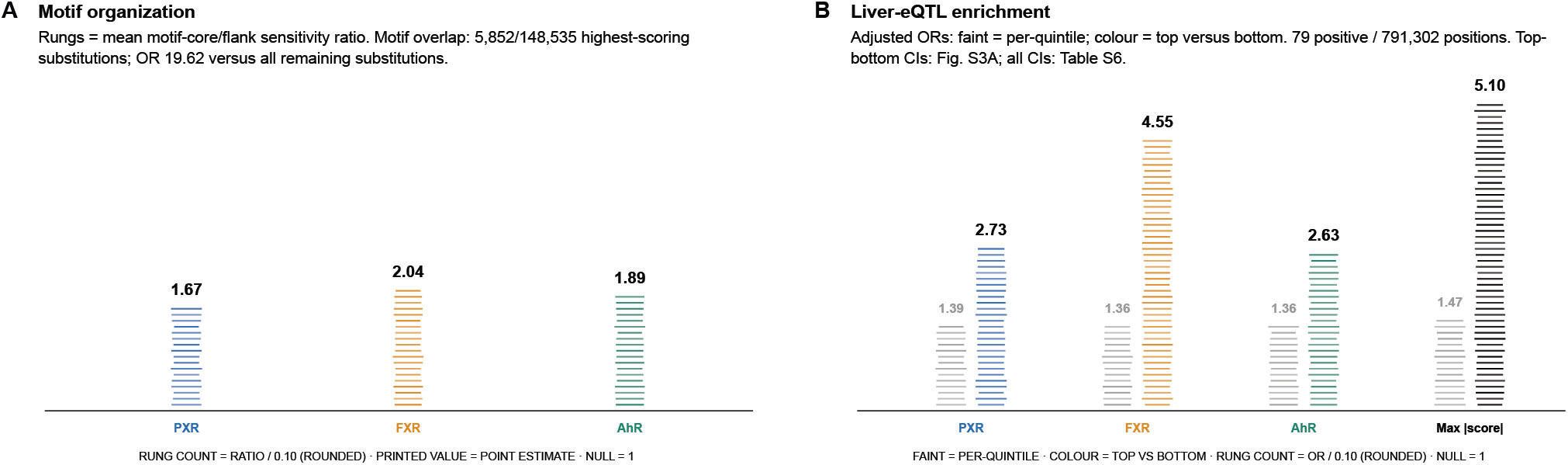
Orthogonal biological organization of the score landscape. (A) Motif core to flank sensitivity ratios and enrichment of substitutions in the top 1% by score within retained motif intervals (OR 19.62). (B) Adjusted odds ratios comparing the top and bottom score quintiles and trends across quintiles among 791,302 positions, including 79 GTEx V10 liver lead eQTL positives. Models adjust for allele frequency and distance to the nearest transcription start site. Confidence intervals are in Supplementary Fig. S3A and Supplementary Table S6. These analyses test biological organization and enrichment, not signed variant effects or clinical outcomes.

Only 79 of 791,302 tested positions were linked to an eQTL. The expression result is therefore interpreted as enrichment rather than validation of effect direction.

All three released reporter alleles with experimentally evaluable directions matched the predicted PXR direction. Standalone scoring recovered the same direction for a fourth case. Because every reporter had informed source selection, no independent validation denominator exists. These cases document functional recovery rather than predictive accuracy. Supplementary Figure S4 and Table S8B provide the records.

## 3.6 Discussion

AetherXeno converts hepatocyte receptor binding into directional scores for every possible substitution in the supported elements. Each score remains linked to evidence for the same allele. The browser, programmatic interface and bulk files expose the same versioned records.

On the test chromosomes, heads trained for each receptor ranked measured occupancy better than the available AlphaGenome and Enformer outputs. The comparison was task specific because the native tracks were receptor or tissue proxies rather than exact experimental matches. These results support adaptation to the studied assays. Comparison with independent models trained for each receptor remains an important next step.

Excluding the four primary windows flagged after repeat masking left the benchmark unchanged, indicating that detectable homology between chromosomes did not account for the observed gains. This audit addresses fitting and testing of the receptor heads, not the AlphaGenome pretraining corpus.

Motif and liver expression analyses independently supported biological organization of the score landscape. Substitutions with high scores concentrated in receptor and cofactor motifs, with greater sensitivity in motif cores than in adjacent flanks. Higher score quintiles were also enriched for liver eQTL positions after adjustment for allele frequency and transcription start site distance. The sparse analysis by genomic position supports prioritization but does not test agreement with the signed expression effect.

Published reporter contrasts provide an interpretable check informed by source selection. All alleles with evaluable directions agreed with the predicted PXR direction, yet those records had contributed to the choice of PXR source input. Prospective reporter testing is therefore required to estimate generalization for individual variants. The *ABCG2* example likewise illustrates evidence use rather than an accuracy endpoint.

Integration by exact allele is distinct from model performance. The release does not transfer expression, clinical or trait evidence to other substitutions at the same base. It also keeps receptor proximity, candidate gene, drug context and literature support separate. The curation layer organizes expert review, but proximity is not proof of the target gene and pharmacist review gates publication. Shared identifiers across the three access routes support reproduction outside the interface.

The atlas currently covers three receptor systems in primary hepatocytes, and the AhR source has one biological replicate. Scores summarize predicted occupancy over 640 bp. They do not measure transcription, drug exposure or clinical response. The benchmark tests ranking of receptor signal against imperfect comparator tracks. The sparse eQTL and reporter analyses provide supporting evidence rather than validation of individual variants. Versioned releases remain essential because external resources change over time.

These constraints suggest concrete extensions without changing the core data model. Additional receptor, tissue and exposure experiments can be added as separate heads when comparable occupancy data and controls are available. Prospective reporter assays can test variants chosen without reference to source selection. Independent models trained for each receptor can broaden the comparison. More extensive expression and perturbation data matched by allele could refine links between occupancy effects and downstream genes. Because AetherXeno retains score definitions, evidence identity, provenance and release versions, such additions can be evaluated against the current resource rather than replacing an untraceable score.

## 3.7 Conclusion

AetherXeno converts receptor occupancy measured in primary human hepatocytes into an exhaustive allele-resolved saturation atlas linked to population, expression, association, clinical and regulatory evidence. Adapting a head for each receptor improved the ranking of measured receptor signal over both comparators on the primary chromosomes. The conclusion was unchanged after conservative homology exclusion. Motif and liver expression analyses showed further biological organization of the score landscape. The browser, programmatic interface and bulk files deliver the same traceable records from individual alleles to cohort workflows while preserving the boundary between predicted occupancy, supporting evidence and clinical interpretation.

## Acknowledgements

The authors thank the investigators who generated and released the primary human hepatocyte datasets used by AetherXeno.

## Author contributions

Y.-H.C.: Conceptualization, data curation, formal analysis, investigation, methodology, project administration, software, validation, visualization, writing—original draft and writing—review and editing. T.-C.C.: Data curation, pharmacological evidence review, validation and writing— review and editing. H.-W.L.: Conceptualization, data curation, validation and writing—review and editing. Y.-C.L.: Conceptualization, project administration, supervision and writing—review and editing.

## Data availability

The AetherXeno browser and application programming interface are publicly accessible at https://aetherxeno.org for noncommercial users without registration. The versioned data release, data dictionary, provenance metadata and checksums are archived in Zenodo record 20753827 at doi:10.5281/zenodo.20753827. Source ChIP-seq data are available under PRJNA239635, GSE57227 and GSE205502. Source code is released under the MIT License. AetherXeno annotations are CC BY-NC 4.0, and outputs derived from AlphaGenome remain subject to the AlphaGenome Output Terms of Use.

