## Supplementary Information for "AetherXeno: an allele-resolved receptor-occupancy atlas for regulatory pharmacogenomics"

Yen-Hung Chen[1], Teng-Che Chuang[2], Hsiang-Wen Lin[3,†] and Yi-Chuan Li[4,*]
[1]Graduate Institute of Biomedical Sciences, China Medical University, Taichung 404328, Taiwan;
[2]China Medical University Hospital, Taichung 404327, Taiwan; [3]School of Pharmacy, China Medical University, Taichung 406040, Taiwan;
[4]Department of Biological Science and Technology, College of Medicine, China Medical University, Taichung 406040, Taiwan

#### Scope

This supplement describes the source experiments, processing, receptor-head adaptation, saturation scoring, evidence integration and evaluation used for the AetherXeno Original Paper. Supplementary Tables S1-S8 contain the complete numerical reporting. AetherXeno scores describe predicted receptor occupancy; they are not calibrated estimates of gene expression, pathogenicity, drug response or treatment effect.

#### Supplementary Methods

##### S1. Source receptor-occupancy experiments

AetherXeno uses public ChIP-seq generated in primary human hepatocytes (PHH). The PXR source was PRJNA239635 (Smith et al., 2014). DMSO- and rifampicin-treated PXR ChIP libraries were pooled for the production target, so the target represents constitutive and ligand-responsive occupancy rather than rifampicin induction alone. The input library served as the peak-calling control.

The FXR source was GSE57227/SRP041597 (Zhan et al., 2014). FXR ChIP-seq followed GW4064 exposure, and the matched rabbit-IgG library was used as background because the study did not provide input DNA.

The AhR source was GSE205502/PRJNA846159 (Filipovic et al., 2023). PHH were exposed to 1 nM TCDD for 24 h. This experiment contained one biological replicate; that limitation is retained in the interpretation of the AhR model. Run-level accessions, controls and read layouts are listed in Supplementary Table S1.

##### S2. Processing and receptor-element definition

Reads were aligned to GRCh38.p13 with BWA-MEM 0.7.19 (Li, 2013). Alignments were coordinate sorted, duplicates were marked and reads with mapping quality below 20 were removed. PXR and AhR single-end coverage was extended to the MACS2-estimated fragment length; FXR coverage used observed paired-end fragments.

Peaks were called with MACS2 2.2.9.1, the study-specific control and one retained duplicate (Zhang et al., 2008). Receptor-specific thresholds and fragment settings are reported in Supplementary Table S2. Because the AhR source had one replicate, its call set was intersected with ENCODE HepG2 ATAC-seq peaks from ENCSR042AWH and ENCFF913MQB (ENCODE Project Consortium 2020). Two prespecified canonical target peaks removed by this filter were restored, producing 3,768 accessibility-supported AhR peaks before final ranking.

Peak summits were ranked within receptor. Overlapping or invalid candidate windows were consolidated, and up to 3,000 elements were retained per receptor. The final release contains 3,000 PXR, 2,947 FXR and 2,744 AhR element records. Their union spans approximately 4.95 Mb of GRCh38 sequence.

##### S3. Receptor-head adaptation and saturation scoring

Fine-tuning used AlphaGenome research implementation v0.2.0 and `google/alphagenome-fold-0` (Avsec et al., 2026). The pretrained sequence trunk was retained with stopped gradients, and one receptor-specific transcription-factor ChIP-seq prediction head was fitted for each receptor. Training examples contained 131,072 bp of sequence and matched receptor coverage at 128-bp resolution. Optimization used Adam; receptor-specific settings and the implementation output key are given in Supplementary Table S2 (Kingma and Ba, 2015).

Chromosomes 2, 8, 11, 15 and 20 were excluded from receptor-head fitting. Five literature-supported PXR regulatory variants on chromosomes 2 and 20 were later consulted when comparing pooled DMSO-plus-rifampicin with rifampicin-only PXR inputs. They were not training labels, but they could influence source-input selection. The primary selection-independent analysis therefore used chromosomes 8, 11 and 15 without refitting or retuning.

At every reference base in the receptor-element union, all three non-reference alleles were generated. For variant v and receptor r, the raw effect summed alternative-minus-reference predictions over the central five 128-bp bins. Receptor-wise p99 scaling was fixed once on the complete receptor-confirmed saturation population. The released score and ranking statistic were defined together as

$$\begin{aligned}
\Delta_r(v) &= \sum_{j=c-2}^{c+2} \left[ f_r(x_v^{\mathrm{ALT}})_j - f_r(x_v^{\mathrm{REF}})_j \right], \\
q_r &= Q_{0.99}\left(\{|\Delta_r(u)| : u \in \mathcal{V}_{\mathrm{receptor-confirmed}}\}\right), \\
\widetilde{\Delta}_r(v) &= \mathrm{clip}\left(\Delta_r(v)/q_r, -1, 1\right), \\
R(v) &= \max_r |\Delta_r(v)|/q_r.
\end{aligned}$$

The five-bin sum covers 640 bp centred on the variant. Positive and negative scores indicate predicted occupancy increase and decrease. Values between $-0.1$ and 0.1 are labelled neutral. The p99 denominators were 38,464 for PXR, 12,608 for FXR and 21,760 for AhR. The normalized score is neither a percentile nor a probability.

The unclipped statistic R was used for cross-receptor ranking. Variants were ordered by decreasing R and then by stable variant identifier. The highest-scoring set comprised the first ceiling of 1% of the receptor-confirmed population. Raw effects, normalized effects and the ranking statistic are retained in the bulk release.

#### S4. Evidence, priority strata and CAMIP

All coordinates and alleles use GRCh38. Population frequencies were obtained from gnomAD genomes v4.1; liver eQTLs from GTEx Analysis Release V10; clinical assertions from the ClinVar GRCh38 file dated 6 June 2026; and associations from the NHGRI-EBI GWAS Catalog snapshot downloaded 12 June 2026 (Chao et al., 2024; GTEx Consortium, 2020; Landrum et al., 2025; Cerezo et al., 2025). GTEx, ClinVar and GWAS joins required exact chromosome-position-reference-alternative identity. Evidence attached to one allele was not propagated to the other substitutions at the same coordinate.

JASPAR 2024.1 CORE vertebrate matrices were scanned with FIMO using a fifth-order Markov background and a nominal threshold of $P<1\times10^{-5}$ (Grant et al., 2011; Rauluseviciute et al., 2024). The receptor/cofactor panel comprised NR1I2/PXR MA1533.2, NR1H4/FXR MA1110.3, AHR::ARNT MA0006.2, RXRA MA0512.2, HNF4A MA0114.5 and FOXA1 MA0148.5. Position-level conservation came from the hg38 phyloP 100-way track (Pollard et al., 2010).

Tier A contains exact-allele GTEx liver-eQTL records. Tier B contains variants in the exact highest-scoring 1% with gnomAD allele frequency below 0.01. These are follow-up strata, not pathogenicity or treatment classes.

CAMIP links a receptor element to a candidate inducible ADME gene when a production peak lies within 25 kb of the transcription start site. The rule generated 29 internal peak-anchored chains: 17 PXR, eight FXR and four AhR. Each record keeps receptor, exposure, locus, predicted consequence, drug context and literature evidence separate, with pharmacogenomic context supported by PharmGKB/ClinPGx or the cited primary literature (Whirl-Carrillo et al., 2021). Pharmacist review gates public release. Proximity does not establish the target gene, and CAMIP does not encode dose, severity, monitoring or prescribing advice.

#### S5. Web application, API and bulk release

The application uses Next.js 16 and React 19. Supabase PostgreSQL serves bounded, read-only queries. The browser and API support variant, rsID, interval, gene, receptor-element, cross-receptor and batch queries; pagination and interval limits prevent unbounded responses. Allele frequency is shown as a percentage in the browser and retained as an exact numeric field in the API and Parquet release.

The service is freely available over HTTPS to non-commercial users without registration at https://aetherxeno.org. Documentation includes examples, a tutorial and field definitions. Versioned Parquet tables, checksums and provenance records are archived at https://doi.org/10.5281/zenodo.20753827 (Chen et al., 2026).

#### S6. Held-out and selection-independent benchmarks

The complete V4 split contained chromosomes 2, 8, 11, 15 and 20. PXR used 1,200 receptor-peak windows and an equal number of accessible receptor-peak-negative windows; FXR and AhR each used 600 peak and 600 background windows. Seventeen 128-bp bins were scored per window.

Identical bins were evaluated with the receptor-specific AetherXeno head, native AlphaGenome and Enformer (Avsec et al., 2021). Native AlphaGenome's HepG2 RXRA track served as the available proxy for PXR and FXR, and its HepG2 AHR track served AhR. Enformer signal was the per-bin mean across one liver CAGE track and three liver H3K27ac tracks, matching the implemented comparator manifest. Measured receptor signal came from the processed ChIP-seq BigWig. Bins overlapping ENCODE HepG2 H3K27ac peaks were excluded from the primary low-baseline stratum.

The primary endpoint was Spearman correlation between predicted and measured signal. AetherXeno-minus-comparator differences were assigned percentile 95% confidence intervals from 1,000 bootstrap samples of whole genomic regions, preserving within-window dependence. Peak-versus-background AUROC was retained only as a secondary diagnostic because both groups were accessible and the contrast was confounded by regional activity.

The selection-independent analysis removed chromosomes 2 and 20, where source-input selection variants occurred, and recomputed the same statistics on chromosomes 8, 11 and 15. Models, predictions, checkpoints and hyperparameters were unchanged. The complete and selection-independent results are reported in Supplementary Tables S3 and S4.

#### S7. Sequence-homology audit

A chromosome holdout prevents coordinate overlap but not duplication across chromosomes. We therefore aligned all 3,534 unique 131,072-bp V4 windows, including 1,764 windows in the selection-independent subset, with minimap2 2.31-r1302 (Li, 2018). The target was a conservative proxy containing every primary GRCh38 chromosome outside chr2, chr8, chr11, chr15 and chr20. Because this proxy includes sequence beyond the actual fine-tuning intervals, it can overestimate potential homology.

Unmasked and RepeatMasker-soft-masked sequences were evaluated at four prespecified length/identity thresholds (Smit et al., 2015). The exclusion analysis removed every window with a repeat-masked alignment of at least 1 kb at 99% identity and then reanalysed the frozen predictions. No model was refitted or retuned. This audit addresses receptor-head fine-tuning/test homology only.

#### S8. Orthogonal and literature-supported analyses

For eQTL enrichment, 871,560 gnomAD-observed SNVs in receptor-confirmed elements were collapsed by genomic position using the maximum absolute score for each receptor, leaving 791,302 positions. GTEx V10 liver significant eQTLs were reduced to the lowest-nominal-P variant for each gene and intersected by position, yielding 79 positives. For each receptor and the maximum-across-receptors axis, the top and bottom score quintiles were compared with

$$\operatorname{logit}\{\Pr(E_i = 1)\} = \beta_0 + \beta_1 T_i + \beta_2 \,\mathrm{MAF}_i + \beta_3 \log_{10}(|d_i| + 1),$$

where E denotes a lead-eQTL position, T the top-quintile indicator and d the distance to the nearest transcription start site. The adjusted odds ratio was $\exp(\beta_1)$. An ordinal-quintile model tested trend; chromosome was added when the sparse-event design remained estimable. This is an enrichment analysis, not validation of the direction or causality of an individual eQTL.

Motif enrichment compared the exact highest-scoring 1% with all remaining receptor-confirmed substitutions by two-sided Fisher's exact test. Local motif grammar was summarized as the ratio of mean per-position maximum absolute raw effect in a retained motif core to the corresponding mean in its adjacent $\pm$50-bp flanks. Core and flank distributions were compared with a one-sided Mann-Whitney U test.

The literature audit considered five PXR regulatory variants. Sugatani et al. (2008; PMID 18172616) tested rs4124874 in a UGT1A1 promoter reporter. Gotoh-Saito et al. (2025; PMID 40301309) tested DPE17 and DPE128 constructs. Factorial DPE128 contrasts isolate rs6013892 and rs158523; a fixed-background DPE17 contrast describes rs3771341; rs4148325 occurs only in multi-variant haplotypes and has no attributable single-allele direction.

All five records had been consulted during PXR source-input selection. Consequently, reporter agreement is reported as supporting recovery and not as independent accuracy. Three exact, direction-evaluable alleles occur in the released atlas; rs3771341 is available only from standalone V1/on-demand scoring, and rs4148325 is indeterminate.

### Supplementary Results

#### S9. Release scale and priority strata

The released atlas contains 17,267,831 unique SNV records with three receptor-effect fields. Of these, 14,853,408 are exhaustive substitutions in receptor-confirmed elements and 3,285,983 have a gnomAD observation; 871,560 records belong to both groups. Exact ranking selected 148,535 highest-scoring substitutions. Tier A contains 1,881 liver-eQTL records, and Tier B contains 6,780 rare high-score records. Complete annotation coverage is given in Supplementary Table S7.

#### S10. Receptor-signal benchmarking

In the secondary complete five-chromosome split, AetherXeno low-baseline Spearman correlations were 0.220, 0.430 and 0.652 for PXR, FXR and AhR. All six AetherXeno-minus-comparator bootstrap intervals excluded zero (Supplementary Table S3).

On the selection-independent chromosomes 8, 11 and 15, the corresponding AetherXeno correlations were 0.211, 0.432 and 0.638. Native AlphaGenome values were 0.117, 0.098 and 0.525; Enformer values were 0.172, 0.328 and 0.458. All six difference intervals again excluded zero (Supplementary Table S4 and Supplementary Figure S1).

#### S11. Homology exclusion sensitivity

Repeat masking reduced the number of windows with a cross-chromosome alignment of at least 1 kb at 99% identity from 65 to 18. None met a longer repeat-masked threshold. Four of the 1,764 selection-independent windows were flagged.

The exclusion removed one PXR, one FXR and two AhR regions from the selection-independent analysis. Rounded correlations were unchanged, the largest absolute correlation change was 0.000232, and all six comparator intervals remained above zero (Supplementary Table S5 and Supplementary Figure S2).

#### S12. eQTL and motif support

Across quintiles of the maximum absolute receptor score, the adjusted trend odds ratio for GTEx liver lead-eQTL enrichment was 1.47 per quintile (95% CI 1.23–1.77; $P=3.21\times10^{-5}$; Main Fig. 4B and Supplementary Table S6). Top-versus-bottom contrasts on each receptor axis are reported in Supplementary Fig. S3A and Supplementary Table S6.

Among the 148,535 highest-scoring substitutions, 5,852 overlapped a retained motif interval. The Fisher odds ratio was 19.62. Mean per-position sensitivity in motif cores exceeded adjacent flanks by 1.673-fold for PXR, 2.040-fold for FXR and 1.885-fold for AhR (Supplementary Fig. S3B and Supplementary Table S8A).

#### S13. Reporter-supported exact alleles

The current release contains three exact PXR reporter alleles with interpretable single-allele directions. Supplementary Fig. S4 focuses on rs6013892 C>A and rs158523 C>T within the same CYP24A1 DPE128 element because their factorial reporter contrasts isolate opposite directions in a shared locus. Their released PXR scores recover those directions. These cases document selection-informed functional recovery rather than an independent accuracy estimate.

Standalone scoring also recovered the positive direction of rs3771341 G>A in a fixed DPE17 background. rs4148325 remained indeterminate because no tested construct isolated its single-allele effect. All four direction-evaluable exact contrasts were concordant descriptively, but no independent accuracy denominator can be estimated because every reporter record had informed source-input selection. Supplementary Fig. S4 shows the released cases; Supplementary Table S8B reports all allele-level values and classifications.

### Supplementary Tables

#### Supplementary Table S1. ChIP-seq source data

| Receptor | Study and run accessions | ChIP condition | Control | Layout; read length | Production target |
|---|---|---|---|---|---|
| PXR | PRJNA239635; SRR1642056 and SRR1642057 | DMSO and rifampicin | SRR1642055 input | Single-end; 36 bp | Pooled DMSO-plus-rifampicin occupancy |
| FXR | GSE57227/ SRP041597; SRR1266979 | GW4064 | SRR1266978 rabbit IgG | Paired-end; 100 bp | Agonist-condition occupancy |
| AhR | GSE205502/ PRJNA846159; SRR19548157 | 1 nM TCDD for 24 h | SRR19548156 input | Single-end; 75 bp | Single-replicate occupancy with ATAC filtering |

#### Supplementary Table S2. Processing, element selection and receptor-head configuration

| Parameter | PXR | FXR | AhR |
|---|---|---|---|
| Aligner / reference | BWA-MEM 0.7.19 / GRCh38.p13 | BWA-MEM 0.7.19 / GRCh38.p13 | BWA-MEM 0.7.19 / GRCh38.p13 |
| Mapping-quality threshold | 20 | 20 | 20 |
| Fragment representation | 138-bp extension | Observed paired fragments | 234-bp extension |
| MACS2 version; q threshold | 2.2.9.1; 0.2 | 2.2.9.1; 0.05 | 2.2.9.1; 0.001 |
| Additional peak filter | Signal rank | Signal rank | HepG2 ATAC ENCFF913MQB plus two-target rescue |
| Selected element records | 3,000 | 2,947 | 2,744 |
| AlphaGenome output | CHIP_TF | CHIP_TF | CHIP_TF |
| Input length | 131,072 bp | 131,072 bp | 131,072 bp |
| Optimizer; learning rate | Adam; $5\times10^{-4}$ | Adam; $5\times10^{-4}$ | Adam; $3\times10^{-4}$ |
| Training steps | 5,000 | 5,000 | 2,000 |
| Global gradient clipping | 0.5 | 0.5 | 0.5 |
| Held-out chromosomes | chr2/8/11/15/20 | chr2/8/11/15/20 | chr2/8/11/15/20 |

#### Supplementary Table S3. Complete five-chromosome V4 benchmark

| Receptor | Bins | Low-baseline bins | AetherXeno $\rho$ | Native AlphaGenome $\rho$ | Enformer $\rho$ | AX-native $\Delta\rho$ (95% CI) | AX-Enformer $\Delta\rho$ (95% CI) |
|---|---|---|---|---|---|---|---|
| PXR | 40,800 | 33,245 | 0.220 | 0.119 | 0.180 | 0.101 (0.086-0.115) | 0.041 (0.028-0.053) |
| FXR | 20,400 | 15,072 | 0.430 | 0.120 | 0.343 | 0.311 (0.271-0.346) | 0.088 (0.062-0.113) |
| AhR | 20,400 | 11,354 | 0.652 | 0.522 | 0.482 | 0.130 (0.113-0.147) | 0.170 (0.139-0.202) |

Values are Spearman correlations in H3K27ac-negative bins. Confidence intervals use 1,000 bootstrap resamples of whole genomic regions.

#### Supplementary Table S4. Selection-independent benchmark on chromosomes 8, 11 and 15

| Receptor | Bins | Low-baseline bins | AetherXeno $\rho$ | Native AlphaGenome $\rho$ | Enformer $\rho$ | AX-native $\Delta\rho$ (95% CI) | AX-Enformer $\Delta\rho$ (95% CI) |
|---|---|---|---|---|---|---|---|
| PXR | 19,720 | 15,718 | 0.211 | 0.117 | 0.172 | 0.094 (0.072-0.115) | 0.039 (0.022-0.056) |
| FXR | 9,384 | 6,835 | 0.432 | 0.098 | 0.328 | 0.334 (0.275-0.392) | 0.104 (0.066-0.145) |
| AhR | 9,979 | 5,081 | 0.638 | 0.525 | 0.458 | 0.113 (0.085-0.138) | 0.180 (0.133-0.226) |

No prediction, model parameter or hyperparameter was changed after chromosomes 2 and 20 were excluded.

#### Supplementary Table S5. Sequence-homology audit and exclusion sensitivity

A. Windows meeting cross-chromosome alignment thresholds

| Analysis population | ≥1 kb / 99% | ≥10 kb / 99% | ≥10 kb / 95% | ≥50 kb / 90% |
|---|---|---|---|---|
| Unmasked, all 3,534 windows | 65 (1.839%) | 3 (0.085%) | 21 (0.594%) | 7 (0.198%) |
| Unmasked, 1,764 chr8/11/15 windows | 21 (1.190%) | 2 (0.113%) | 10 (0.567%) | 1 (0.057%) |
| Repeat-masked, all 3,534 windows | 18 (0.509%) | 0 | 0 | 0 |
| Repeat-masked, 1,764 chr8/11/15 windows | 4 (0.227%) | 0 | 0 | 0 |

B. Selection-independent results after excluding repeat-masked ≥1 kb / 99% windows

| Receptor | Regions excluded | Bins / low-baseline bins | AetherXeno $\rho$ | Native AlphaGenome $\rho$ | Enformer $\rho$ | AX-native $\Delta\rho$ (95% CI) | AX-Enformer $\Delta\rho$ (95% CI) |
|---|---|---|---|---|---|---|---|
| PXR | 1 | 19,703 / 15,701 | 0.211 | 0.117 | 0.172 | 0.094 (0.073-0.115) | 0.039 (0.022-0.058) |
| FXR | 1 | 9,367 / 6,835 | 0.432 | 0.098 | 0.328 | 0.334 (0.275-0.392) | 0.104 (0.066-0.145) |
| AhR | 2 | 9,945 / 5,080 | 0.638 | 0.525 | 0.458 | 0.113 (0.085-0.139) | 0.181 (0.133-0.224) |

#### Supplementary Table S6. GTEx V10 liver eQTL enrichment

| Score axis | Top/bottom positions | eQTLs in top / bottom | Adjusted OR (95% CI) | P | OR per quintile (95% CI) | Trend P | Chromosome sensitivity |
|---|---|---|---|---|---|---|---|
| PXR | 318,775 | 28 / 9 | 2.73 (1.24-5.99) | 0.0123 | 1.39 (1.17-1.66) | $2.23\times10^{-4}$ | Not estimable (singular design) |
| FXR | 335,691 | 28 / 7 | 4.55 (1.96-10.56) | $4.20\times10^{-4}$ | 1.36 (1.14-1.61) | $4.91\times10^{-4}$ | 4.81 (2.06-11.23), P=$2.91\times10^{-4}$ |
| AhR | 324,081 | 31 / 12 | 2.63 (1.29-5.36) | 0.00808 | 1.36 (1.14-1.62) | $5.86\times10^{-4}$ | 2.60 (1.25-5.39), P=0.0103 |
| Maximum across receptors | 314,370 | 32 / 6 | 5.10 (2.01-12.96) | $6.09\times10^{-4}$ | 1.47 (1.23-1.77) | $3.21\times10^{-5}$ | Not estimable (singular design) |

The primary model included all 791,302 positions and 79 eQTL-positive positions before top/bottom restriction and adjusted for MAF and $\log_{10}(|\text{distance to TSS}|+1)$.

#### Supplementary Table S7. Release versions, scale and score parameters

A. Reference and annotation versions

| Resource | Release/file used |
|---|---|
| Genome | GRCh38.p13 |
| AlphaGenome | Research implementation v0.2.0; `google/alphagenome-fold-0` |
| ENCODE accessibility | HepG2 ATAC ENCSR042AWH / ENCFF913MQB |
| ENCODE baseline chromatin | HepG2 H3K27ac ENCSR000AMO / ENCFF580KMC |
| gnomAD | Genomes v4.1 sites files |
| GTEx | Analysis Release V10; liver significant-pairs Parquet |
| ClinVar | GRCh38 VCF, file date 6 June 2026 |
| GWAS Catalog | Associations snapshot downloaded 12 June 2026 |
| JASPAR | 2024.1 CORE vertebrates |
| Conservation | hg38 phyloP 100-way, retrieved 19 June 2026 |
| PharmGKB/ClinPGx | Downloadable archive retrieved 4 June 2026 |

B. Release scale and annotation coverage

| Component | Count |
|---|---|
| Unique scored SNV records | 17,267,831 |
| Exhaustive receptor-confirmed substitutions | 14,853,408 |
| gnomAD-observed released records | 3,285,983 |
| Receptor-element records | 8,691 |
| Allele-matched GTEx liver-eQTL records / Tier A | 1,881 |
| Tier A records linked to the curated ADME panel | 5 |
| ClinVar records | 28,227 |
| Allele-matched GWAS Catalog associations | 856 |
| Receptor-confirmed substitutions overlapping a retained JASPAR/FIMO motif interval | 36,528 |
| phyloP-annotated receptor-confirmed substitutions | 14,804,721 |
| Exact highest-scoring 1% | 148,535 |
| Tier B high-score rare substitutions | 6,780 |
| Internal CAMIP peak-anchored chains | 29 |

##### C. Global score parameters

| Parameter | PXR | FXR | AhR |
|---|---|---|---|
| p99 absolute raw-effect denominator | 38,464 | 12,608 | 21,760 |
| Released display range | −1 to 1 | −1 to 1 | −1 to 1 |
| Decrease / neutral / increase | <−0.1 / [−0.1, 0.1] / >0.1 | <−0.1 / [−0.1, 0.1] / >0.1 | <−0.1 / [−0.1, 0.1] / >0.1 |

#### Supplementary Table S8. Orthogonal biological support

##### A. Target-motif core versus adjacent flank sensitivity

| Receptor | Core positions | Flank positions | Mean core sensitivity | Mean flank sensitivity | Core/flank ratio | One-sided Mann-Whitney P |
|---|---|---|---|---|---|---|
| PXR | 5,260 | 35,002 | 19,196.08 | 11,476.22 | 1.673 | $4.30\times10^{-49}$ |
| FXR | 4,862 | 36,666 | 12,513.65 | 6,132.99 | 2.040 | $<10^{-300}$ |
| AhR | 3,062 | 25,593 | 13,905.95 | 7,376.21 | 1.885 | $8.60\times10^{-160}$ |

##### B. PXR reporter records used for supporting recovery

| Variant | GRCh38 allele | Experimental interpretation | Released PXR raw Δ / normalized | V1 raw Δ / validation-background percentile | Evaluation classification |
|---|---|---|---|---|---|
| rs4124874 | chr2:233757013 T>G | T>G reduced PXR/ rifampicin-responsive UGT1A1 reporter activation (PMID 18172616) | −7,744 / −0.201331 | −8,000 / 93.66 | Released exact allele; direction-concordant |
| rs3771341 | chr2:233764593 G>A | Fixed DPE17 background, A versus G increased reporter activity by 122.5% (PMID 40301309) | Not released | +6,464 / 92.46 | On-demand exact contrast; direction-concordant |
| rs4148325 | chr2:233764663 C>T | Tested only in multi-variant DPE17 haplotypes (PMID 40301309) | Not released | +768 / 66.15 | Single-allele direction indeterminate |
| rs6013892 | chr20:54107677 C>A | DPE128 factorial contrasts −43.4% and −42.3% (PMID 40301309) | −25,984 / −0.675541 | −26,496 / 98.33 | Released exact allele; direction-concordant |
| rs158523 | chr20:54107770 C>T | DPE128 factorial contrasts +62.9% and +59.9% (PMID 40301309) | +5,376 / +0.139767 | +5,376 / 90.67 | Released exact allele; direction-concordant |

V1 percentiles are ranks against a fixed validation background, not percentiles in the public release. All five rows are source-selected supporting cases and contribute zero observations to an independent accuracy denominator.

#### Supplementary Figure Captions

#### Supplementary Figure S1. Selection-independent receptor-signal benchmark details

(A) Spearman correlations between predicted and measured receptor ChIP signal in H3K27ac-negative bins on chromosomes 8, 11 and 15. Identical bins were evaluated with AetherXeno, native AlphaGenome and Enformer. (B) AetherXeno-minus-comparator differences in Spearman correlation. Points are observed differences; intervals are percentile 95% confidence intervals from 1,000 region-block bootstrap resamples. (C) Held-out chromosomes and the tested-bin, low-baseline-bin and bootstrap-region denominators for each receptor. Exact values and intervals are reported in Supplementary Table S4.

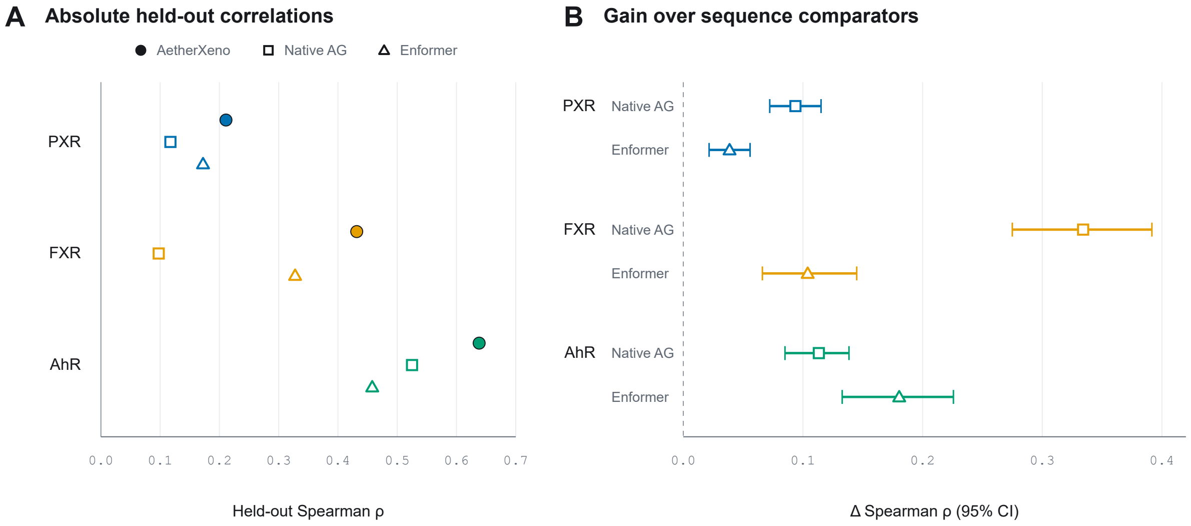

#### Supplementary Figure S2. Cross-chromosome homology audit

(A) Counts of 3,534 held-out windows meeting four cross-chromosome alignment thresholds before and after RepeatMasker soft masking: 65 to 18, 3 to 0, 21 to 0 and 7 to 0. (B) Change in selection-independent low-baseline Spearman correlation after exclusion of repeat-masked windows with alignments ≥1 kb at ≥99% identity, shown as post-exclusion minus original values ($\times 10^{-4}$) for each receptor and model. The largest unscaled absolute change was 0.000232. This audit addresses receptor-head fine-tuning/test separation and does not characterize AlphaGenome pretraining.

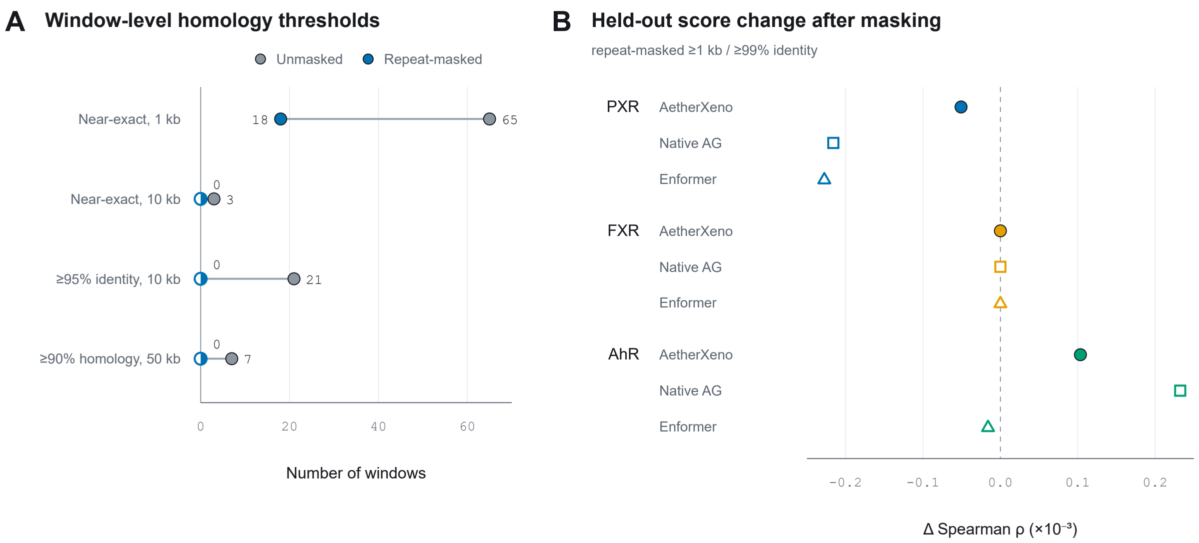

#### Supplementary Figure S3. Orthogonal biological support

(A) Adjusted odds ratios for GTEx V10 liver lead-eQTL enrichment in the top versus bottom score quintiles among 791,302 tested positions, including 79 eQTL-positive positions. Models adjust for MAF and log10(|distance to TSS|+1); intervals are Wald 95% confidence intervals. (B) Ratio of mean absolute per-position saturation sensitivity in retained target-motif cores to adjacent ±50 bp flanks. The reference line denotes a ratio of one. These analyses support positional and motif-level biological prioritization; they do not validate signed SNV effects or clinical outcomes.

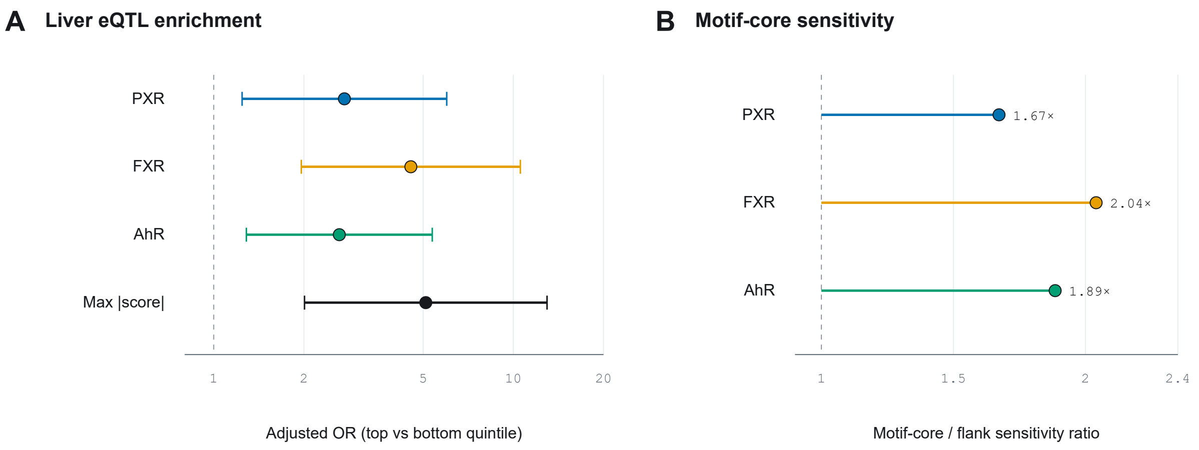

#### Supplementary Figure S4. Descriptive exact-allele concordance

Shared-locus comparison of rs6013892 C>A and rs158523 C>T in the CYP24A1 DPE128 PXR element. The two variants have opposite released PXR score directions and opposite factorial reporter effects reported by Gotoh-Saito et al. (2025). The cases informed PXR source-input selection and therefore document supporting functional recovery rather than an independent predictive test. The complete allele audit is reported in Supplementary Table S8B.

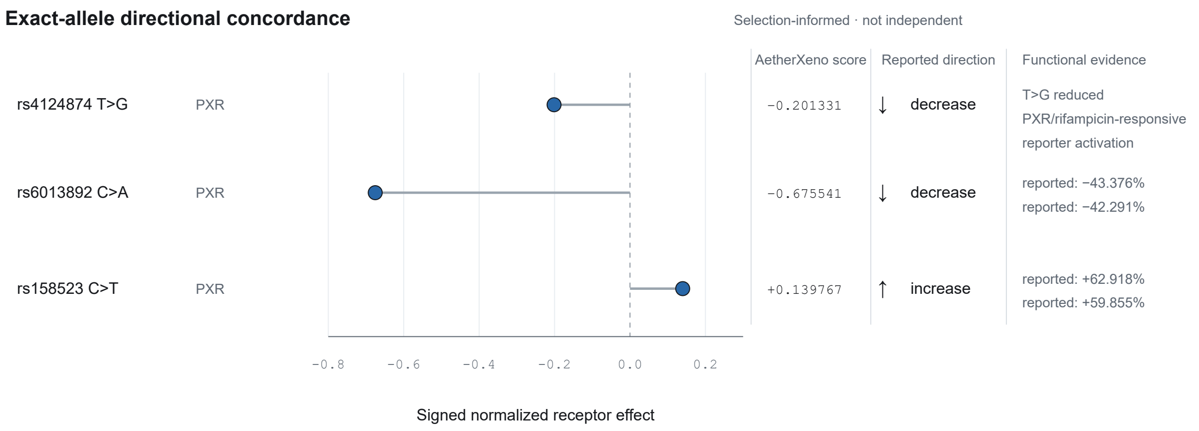
